# Nitrate-regulated growth processes involve activation of gibberellin pathway

**DOI:** 10.1101/2021.07.27.453969

**Authors:** Lucie Camut, Barbora Gallova, Lucas Jilli, Mathilde Sirlin-Josserand, Esther Carrera, Lali Sakvarelidze-Achard, Julie Zumsteg, Dimitri Heintz, Sandrine Ruffel, Gabriel Krouk, Stephen G. Thomas, Peter Hedden, Andrew L. Phillips, Jean-Michel Davière, Patrick Achard

**Author notes:** Correspondence: Dr Patrick Achard, Institut de biologie moléculaire des plantes du CNRS, 12, rue Général Zimmer, 67084 Strasbourg Cedex, France.

## Abstract

Nitrate, one of the main nitrogen (N) sources for crops, acts as a nutrient and key signaling molecule coordinating gene expression, metabolism and various growth processes throughout the plant life cycle. It is widely accepted that nitrate-triggered developmental programs cooperate with hormone synthesis and transport, to finely adapt plant architecture to N availability. Here, we report that nitrate, acting through its signaling pathway, promotes growth in *Arabidopsis* and wheat, in part by modulating the accumulation of gibberellin (GA)-regulated DELLA growth repressors. We show that nitrate reduces the abundance of DELLAs by increasing GA contents through activation of GA metabolism gene expression. Consistently, the growth restraint conferred by nitrate deficiency is partially rescued in *global-DELLA* mutant that lacks all DELLAs. At the cellular level, we show that nitrate enhances both cell proliferation and elongation in a DELLA-dependent and -independent manner, respectively. Our findings establish a connection between nitrate and GA signaling pathways that allow plants to adapt their growth to nitrate availability.

## Introduction

Nitrogen (N) is one of the most important macronutrients limiting plant growth and yield [1-3]. Except for a few plant species that can use atmospheric N_2_ gas through symbiotic association with certain soil bacteria, most crops acquire N in the soil from inorganic forms such as nitrate (NO_3_^-^), the main N source for plants in agricultural lands [4]. Nitrate serves as a source of the essential element found in very diverse macromolecules and a signal molecule regulating global gene expression and modulating both root and shoot system architecture [2, 3, 5-7].

The most studied nitrate-related signaling pathway is called the primary nitrate response (PNR) [8]. It is characterized by a fast nitrate specific (manifested in *nitrate reductase*-null mutant; [5]) induction of marker genes (reviewed in [6]). These markers include the nitrate transporter families such as the transceptor *NITRATE TRANSPORTER 1 (NRT1)/PEPTIDE TRANSPORTER (PTR) FAMILY 6*.*3* (*NPF6*.*3*, also known as *CHLORATE RESISTANT 1, CHL1* or *NRT1*.*1*), *NRT2*.*1* and *NRT2*.*2*, the nitrate assimilation genes *NITRITE REDUCTASE* (*NiR*) and *NITRATE REDUCTASE* (*NIA1* and *NIA2*) and a multitude of transcription factors functioning in diverse signaling pathways [2, 7, 9-12]. According to the current model of PNR, NRT1.1 perceives nitrate triggering changes in cytoplasmic Ca^2+^ levels [13], which activate several members of calcium-sensor protein kinases (CPK10/30/32; [14]) that phosphorylate conserved NIN-LIKE PROTEIN (NLP) transcription factors (NLP6/7; [15]) to reprogram nitrate-regulated genes ensuring adaptive growth to nitrate availability [14, 16].

Despite sophisticated nitrate uptake, storage and assimilation mechanisms [12], the fluctuation of nitrate concentration in both time and space, in part due to its high solubility and predisposition to leaching, requires farmers to supply soil with N-fertilizers to ensure optimal crop yield. Hence, since the 1960s, the application of N-fertilizers associated with mechanization and the adoption of high-yielding semi-dwarf varieties led to substantial yield increases, an intensive agricultural practice known as the “Green Revolution” [17]. The resulting semi-dwarf stature prevents lodging and thus enables higher N-fertilizer applications. The semi-dwarfing genes of the rice (*semidwarf1, sd1*) and wheat (*Reduced height 1, Rht-1*) varieties were characterized and shown to inhibit the synthesis and action, respectively, of gibberellin (GA) growth hormones [17]. Strikingly, despite that more than 70% of wheat cultivated worldwide carry a GA-insensitive *Rht-1* dwarfing allele [18], little is known about whether and how GA contributes to nitrate-regulated growth processes.

GAs constitute a class of diterpenoid molecules controlling major aspects of plant growth and development. Mutants deficient in GA biosynthesis or responses are dwarfs and late flowering, while elevated GA signaling results in taller plants and early flowering. Bioactive GA promotes growth by opposing the functions of DELLA growth repressing proteins (DELLAs), members of the GRAS family of transcriptional regulators [19]. While cereals usually harbor a single DELLA paralogue, such as *SLENDER RICE 1* (*SLR1*) in rice and *REDUCED HEIGHT 1* (*RHT-1*) in wheat [20], the *Arabidopsis* genome encodes five DELLAs, GA-INSENSITIVE (GAI), REPRESSOR of *ga1-3* (RGA), RGA-LIKE1 (RGL1), RGL2 and RGL3 [19]. GA-mediated physiological responses are activated by the binding of bioactive GA to the GA receptors GIBBERELLIN INSENSITIVE 1 (GID1) that trigger the destruction of DELLAs through the ubiquitin-dependent 26S proteasome pathway [19]. DELLAs repress GA responses by interacting with and modulating the activity of DELLA-interacting partners (DIP) such as transcription factors or regulators, and chromatin-remodeling complexes [21]. Thereby, DELLAs regulate the expression of a broad array of genes involved in various pathways.

Similar to other phytohormones, GAs act as mediators of environmental signals, allowing plants to respond, often rapidly, to changes in light conditions, temperature, water and nutrient status [22-26]. Usually, DELLA accumulation reduces growth to prioritize resources to defense mechanisms, whereas GA-mediated DELLA degradation stimulates growth under favorable conditions [27]. Hence, it is now accepted that GA signaling enables appropriate growth in adverse environments [22, 27]. Interestingly, recent experiments have shown that physical interactions between the rice DELLA SLR1 and GROWTH-REGULATOR FACTOR 4 (GRF4) transcription factor co-regulate growth and the metabolism of carbon and nitrogen [28]. Whereas GRF4 promotes N assimilation, carbon fixation and plant growth, DELLAs repress these processes. Moreover, a recent study has reported that SLR1, and also GID1, are able to interact with NITROGEN-MEDIATED TILLER GROWTH RESPONSE 5 (NGR5), a rice APETALA2-domain transcription factor induced by N [29]. SLR1 accumulation competitively inhibits GID1-NGR5 interaction, which in turn enhances NGR5 stability and thereby shoots branching [29]. Thus, although it is apparent that GA signaling regulates growth and nitrogen-use efficiency, at present, it is currently unclear whether GA is a component of the regulatory pathway controlling the nitrate-dependent growth. Here, we undertook a molecular and genetic approach to evaluate the role of GA metabolism and signaling in nitrate-regulated growth processes in *Arabidopsis* and wheat. Essentially, we show that nitrate increases bioactive GA levels, thus promoting the degradation of DELLAs. Reduced DELLA accumulation in turn activates cell proliferation, and as a consequence, root and shoot growth. Plant growth adaptation to fluctuating nutrient environment can be understood as a feed-forward cycle, where nutrients promote plant growth through regulating hormones biosynthesis and transport, and in turn, hormonal pathways control nutrient provision to growth rate [30, 31]. Our data lead to the proposal that the GA pathway is a major regulator of this feed-forward loop interconnecting plant growth to nitrate availability.

## Results

### Nitrate regulates plant growth in a DELLA-dependent manner

In response to nitrate availability, the root and shoot system architecture undergoes important developmental changes. Thus, while adequate nitrate availability increases the length of both primary and lateral roots, low and high nitrate supplies inhibit their growth [32]. Similarly, nitrate limitation reduces shoot growth and delays axillary bud activation and therefore shoot branching [33]. Although well described as growth regulators preventing excessive elongation in response to high N fertilizer supply, it is still unclear whether the GA-regulated DELLA proteins inhibit growth under low N regimes [17, 18]. To assess this point, we first investigated the involvement of DELLAs in the regulation of primary root growth in the absence of nitrate (or 0.5 mM glutamine, an alternative N source) and in response to increasing amounts of nitrate. For this purpose, we compared the primary root length of *Arabidopsis* wild-type and *global-DELLA* mutant seedlings (lacking all five *Arabidopsis* DELLAs) grown with nitrate supply ranging from 0 to 50 mM (Fig. 1A, B). As previously reported [32], the growth of the primary roots was proportionally correlated with the amount of nitrate, from low to adequate concentrations (10 mM NO_3_^-^ in our growth conditions), and then inversely correlated under high nitrate supply (50 mM NO_3_^-^). Interestingly, we observed that the primary root growth of 7-d-old *global DELLA* mutant seedlings was less affected by low and high nitrate supply than that of the wild-type (Fig. 1B). Whereas the length of *global-DELLA* mutant seedling roots grown on 0.25, 0.5, 1 and 50 mM NO_3_^-^ was significantly longer than wild-type roots, their differences decreased on adequate nitrate supply (10 mM NO_3_^-^). Thus, DELLA function inhibits root growth in response to low and high nitrate availability, but becomes less critical on optimal nitrate supply (Fig. 1A, B).

**Figure 1.**
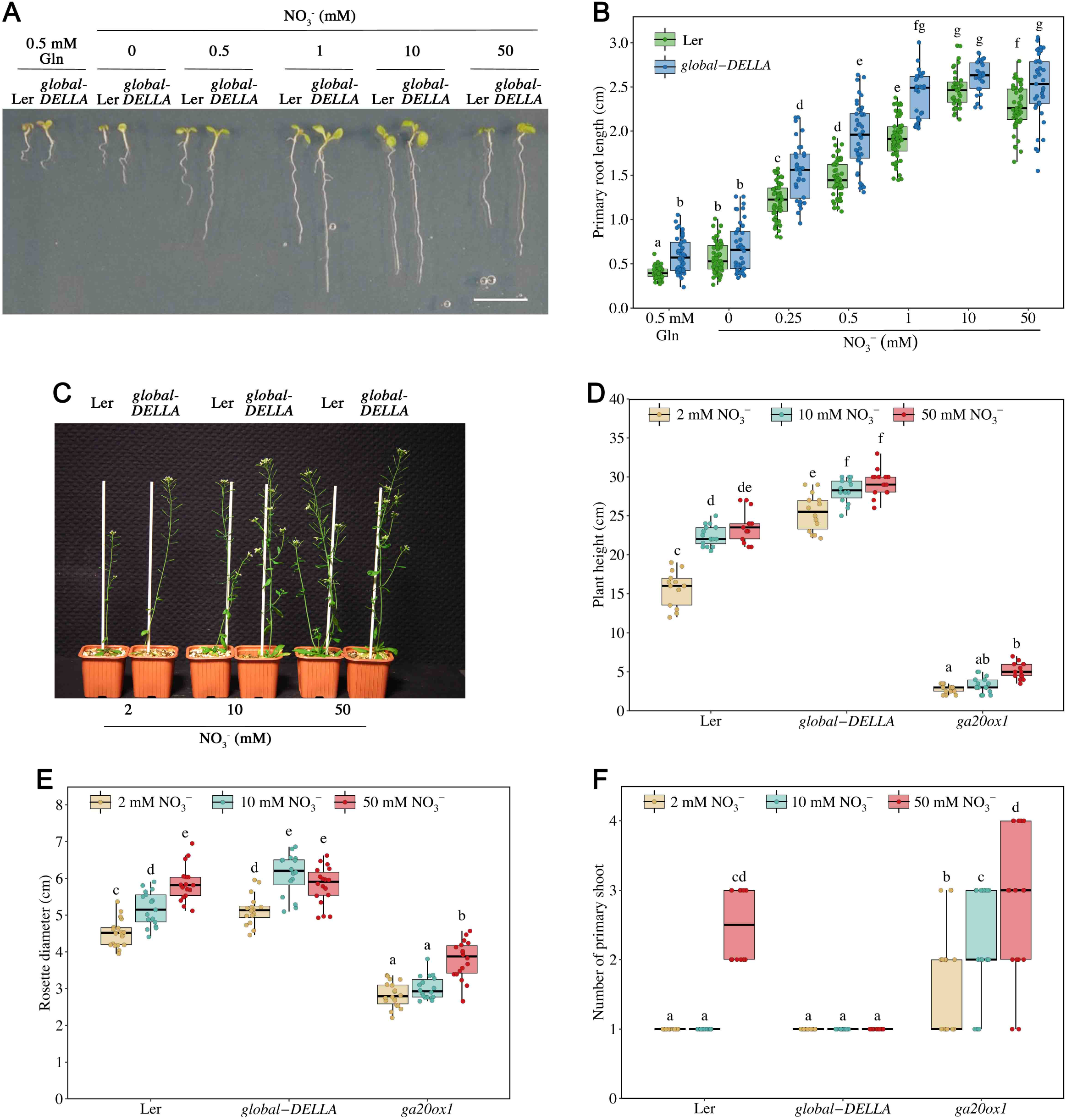
Nitrate regulates plant growth in part in a DELLA-dependent manner. **A**, Representative 7-day-old wild-type (Ler) and *global DELLA* mutant seedlings grown on media containing increasing concentrations of nitrate (NO_3_^-^), as indicated. Gln, glutamine. Scale bar represents 1 cm. **B**, Root length of 7-day-old wild-type (Ler) and *global DELLA* mutant seedlings grown on increasing concentration of nitrate, measured between day 2 and day 7 post germination. Different letters denote significant differences (*p* < 0.05) from the wild-type using two-way ANOVA followed by a Tukey’s test for multiple comparisons. 25 ≤ n ≤ 61 seedlings were analyzed for each box plots. Similar results were obtained in independent experiments. **C**, Representative 7-week-old wild-type (Ler) and *global DELLA* mutant plants grown on perlite vermiculite soil supplemented once a week with a nutritive solution containing 2, 10 or 50 mM NO_3_^-^. **D-F**, Height of 7-week-old (13 ≤ n ≤ 17 plants) (**D**), rosette diameter of 5-week-old (16 ≤ n ≤ 18 plants) (**E**) and number of primary shoot of 8-week-old (14 ≤ n ≤ 19 plants) (**F**) wild-type (Ler), *global DELLA* and *ga20ox1* mutant plants grown on different nitrate conditions, such as in **C**. Different letters denote significant differences (*p* < 0.05) using two-way ANOVA with Tukey’s test for multiple comparisons. Similar results were obtained in independent experiments.

Next, to examine the contribution of DELLAs to nitrate-mediated growth processes during shoot development, we grew *Arabidopsis* wild-type, *global DELLA* and *ga20ox1* mutants (a weak GA biosynthesis mutant [34-36]) on a perlite vermiculite soil supplemented with increasing amounts of nitrate (ranging from 2 to 50 mM NO_3_^-^). It is noteworthy that nitrate concentrations used for *in vitro* experiments (Fig. 1A, B) may not be directly comparable to those applied in soil. Nevertheless, in a similar way to the above growth responses, we found that whereas the height of 7-week-old wild-type plants increased with the concentration of nitrate present in the soil, the growth response was attenuated in mutants altered in GA pathway (Fig. 1C, D). Furthermore, while the diameter of wild-type rosettes became larger when nitrate supply was increased, the *global-DELLA* mutant rosettes were insensitive to a change from 10 to 50 mM NO_3_^-^ (Fig. 1E). Taken together, these results demonstrate that nitrate promotes growth in part via a DELLA-dependent mechanism.

There is considerable evidence suggesting that nitrate-dependent regulation of bud activation and elongation are triggered by changes in hormone action [33]. GA has long been known to repress shoot branching; GA-deficient mutants display a high degree of branching [37]. We found that nitrate supply increased the number of primary branches of wild-type and *ga20ox1* mutant plants in contrast to *global-DELLA* mutant plants (Fig. 1F). Thus, similar to the above results, nitrate enhances bud outgrowth in a DELLA-dependent manner. At this stage, it remains unclear whether the NGR5-GID1-DELLA competitive binding mechanism controlling shoot branching in rice also operates in *Arabidopsis* [29].

### Nitrate promotes root meristem growth via DELLA-dependent action on cell proliferation

Plant growth is regulated through the activity of apical and intercalary meristems in which the cells divide before elongating and differentiating. Moreover, GA promotes growth through cell proliferation and cell elongation by stimulating the degradation of nuclear DELLAs [19, 38]. Previous works reported a key role for DELLAs in controlling root apical meristem (RAM) activity [39, 40]. To investigate whether DELLAs play an essential function in regulating RAM activity in response to nitrate availability, we measured the size of the RAM of *Arabidopsis* wild-type and *global-DELLA* mutant seedlings throughout their development (from 3 to 7-d-old) on low (0.5 mM) and adequate (10 mM) NO_3_^-^ conditions. Root meristem length was represented as the number of cortical cells between the quiescent center (QC) and the first elongated cell [40]. We found that whereas the size of the root meristem of wild-type seedlings grown on 10 mM NO_3_^-^ was significantly increased compared to those grown on 0.5 mM NO_3_^-^, the *global-DELLA* mutant root meristem was less sensitive to the effect of nitrate (Fig. 2A).

**Figure 2.**
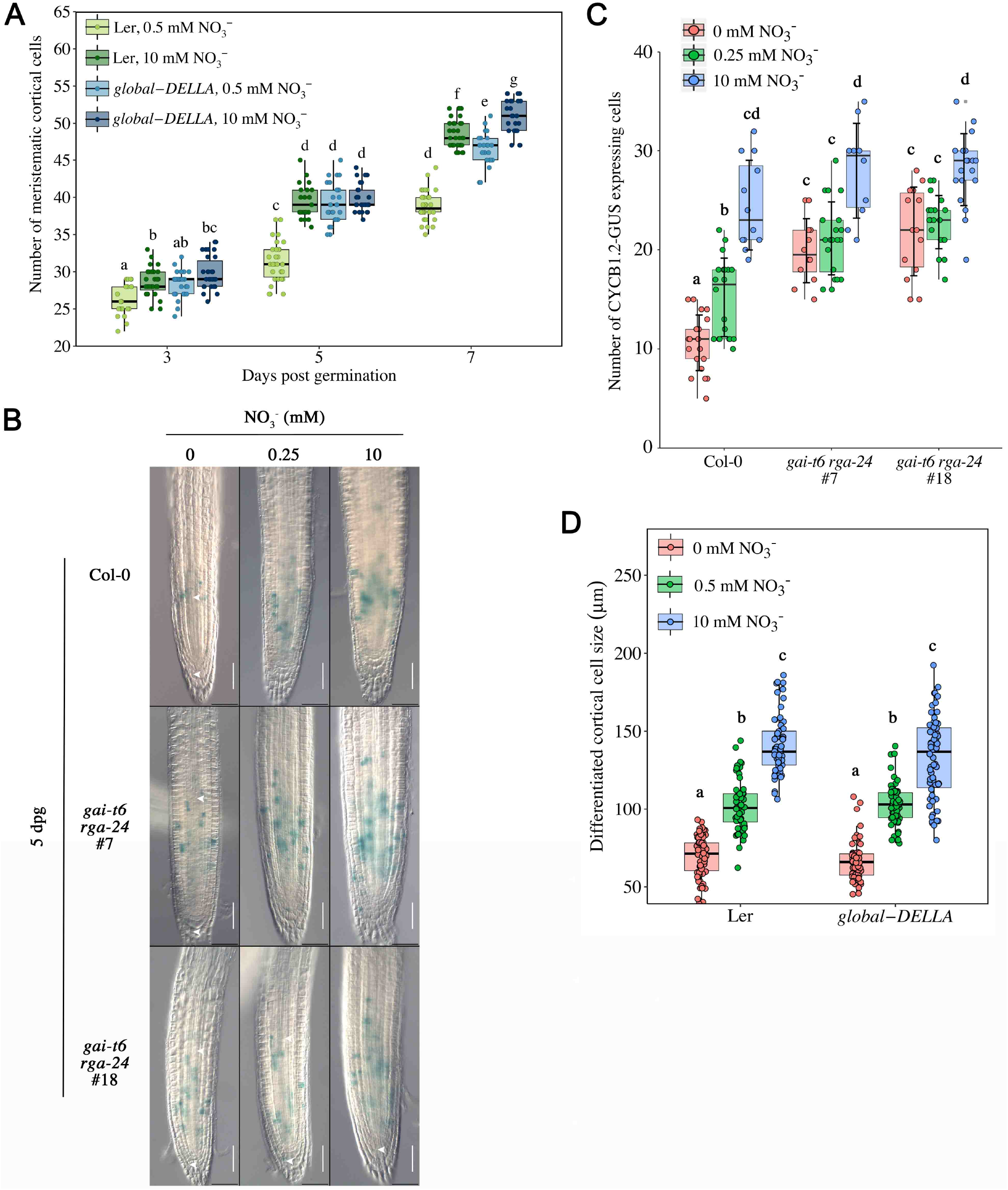
Nitrate promotes root meristem cell division in a DELLA-dependent manner. **A**, Root meristem cell number of wild-type (Ler) and *global DELLA* mutant seedlings at 3, 5 and 7 days post germination (dpg), grown on 0.5 or 10 mM NO_3_^-^. The number of meristematic cells was determined in the cortex file between the quiescent center and the first differentiated cells. Different letters denote significant differences (*p* < 0.05) using two-way ANOVA with Tukey’s test for multiple comparisons. 20 ≤ n ≤ 30 cortex files from at least 11 roots were analyzed for each box plot. Similar results were obtained in independent experiments. **B-C**, Effect of NO_3_^-^ treatment on CYCB1;2:Dbox-GUS cell-division marker in wild-type (Col-0) and *gai-t6 rga-24* mutant (lines #7 and #18) roots. (**B**) Photographs show representative 5-day-old root seedlings grown without nitrate (0 mM) or with 0.25 or 10 mM NO_3_^-^. White arrows indicate the distance between the first and last dividing cells in the longitudinal axis of the root meristem as shown by ß-glucuronidase staining. Scale bars represent 100 µm. (**C**) Number of CYCB1;2:Dbox-GUS expressing cells in root meristems of 5-d-old seedlings, as in **B**. Different letters denote significant differences (*p* < 0.05) using one-way ANOVA with Tukey’s test for multiple comparisons. 10 ≤ n ≤ 21 roots were analyzed for each box plot. **D**, Length of differentiated cortical cells of 7-day-old wild-type (Ler) and *global DELLA* mutant seedlings grown without nitrate (0 mM) or with 0.5 or 10 mM NO_3_^-^. Different letters denote significant differences (*p* < 0.05) using two-way ANOVA with Tukey’s test for multiple comparisons. 51 ≤ n ≤ 66 differentiated cortical cells were analyzed for each box plot. Similar results were obtained in independent experiments.

To corroborate this finding, we also analyzed the activity of the mitotic CYCLIN B1;2-GUS reporter in wild-type and *gai-t6 rga-24* double mutant (a mutant lacking the two main DELLAs restraining growth) [40]. We observed that the number of dividing cells significantly decreased in the RAM of wild-type seedlings grown in nitrate deficient condition compared with those grown on 10 mM NO_3_^-^ (Fig. 2B, C). In contrast, the number of dividing cells in the *gai-t6 rga-24* RAM was less affected by the reduction in nitrate availability. Taken together, these results indicate that DELLAs contribute to the nitrate-induced cell division activity.

Because root growth is the result of both cell division and cell expansion, we next asked whether nitrate and DELLAs jointly regulate root cell elongation. To this end, we measured the length of differentiated cortical cells in the root differentiation zone of 7-d-old wild-type and *global-DELLA* mutant seedlings grown on low (0.5 mM) and adequate (10 mM) NO_3_^-^ conditions. We found that nitrate increased the length of the cells at a similar level in wild-type and *global-DELLA* mutant seedling roots (Fig. 2D). Thus despite reports that GA/DELLA regulates cell elongation in root [38], adequate nitrate supply (10 mM NO_3_^-^) enhances cell expansion in a DELLA-independent fashion.

### Low nitrate availability enhances DELLA protein accumulation

In a subsequent experiment, we investigated whether the nitrate-regulated growth processes are associated with DELLA protein accumulation. To this end, we used *pRGA:GFP-RGA* transgenic seedlings expressing a functional GFP-RGA protein visible in nucleus of root cells [41]. We found a significant decrease of the GFP-RGA signal in the RAM of seedlings grown on 1 and 10 mM NO_3_^-^ compared to those grown on low nitrate conditions (Fig. 3A). By contrast, GFP-RGA signal was increased in RAM of seedlings grown on Gln or ammonium chloride (NH_4_Cl), two alternative N sources, thus confirming that this response is specific to nitrate (Fig. S1). Remarkably, consistent with the above growth phenotypes, we also observed an increase in GFP-RGA signal in seedling roots grown on high nitrate conditions (50 mM NO_3_^-^; Fig. 3A). Then, we investigated the effect of nitrate supply on GFP-RGA protein accumulation in seedling roots and shoots. Accordingly, we found that nitrate substantially decreases the accumulation levels of GFP-RGA protein in both roots and shoots, but their sensitivity to nitrate was different. Indeed, while 1 mM NO_3_^-^ was sufficient to trigger a significant reduction in GFP-RGA level in roots, 10 mM NO_3_^-^ was required to induce the same response in shoots (Fig. 3B, C and independent replicate shown in Fig. S2A, B). Finally, to further substantiate the effect of nitrate on DELLA protein abundance, we determined the accumulation levels of endogenous RGA protein in shoots and roots of seedlings transferred from nitrate deficiency to adequate nitrate conditions (10 mM NO_3_^-^), and inversely. Time-course analysis in seedling shoots revealed that nitrate supply decreases RGA protein abundance and nitrate deficiency increases RGA accumulation within 48 hr (Fig. 3D, E and independent replicate shown in Fig. S2C, D). A similar response was also observed in seedling roots, although the difference was less pronounced (Fig. 3F, G and independent replicate shown in Fig. S2E, F).

**Figure 3.**
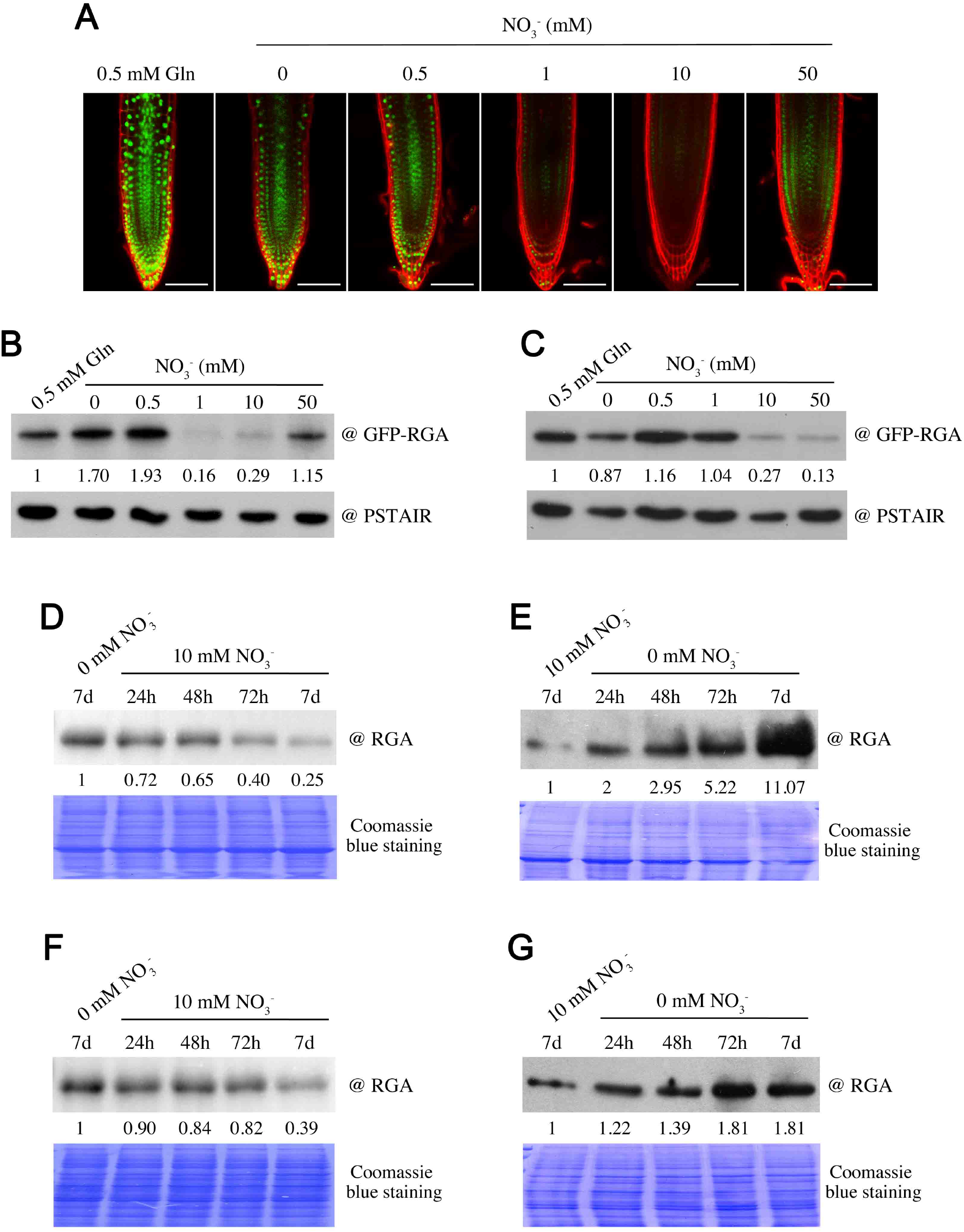
NO_3_^-^-regulated growth processes are associated with DELLA accumulation. **A**, GFP fluorescence in root meristem of 7-day-old *pRGA:GFP-RGA* seedlings grown on media containing increasing concentration of nitrate (NO_3_^-^), as indicated. All images were obtained with the same parameters. Gln, glutamine. Scale bars represent 100 µm. **B-C**, Immunodetection of GFP-RGA (with an antibody to GFP) in roots (**B**) and shoots (**C**) of 7-day-old *pRGA:GFP-RGA* seedlings grown on media containing increasing concentration of NO_3_^-^, such as in **A**. PSTAIR serves as sample loading control. Numbers represent the fold increase in GFP-RGA protein levels relative to PSTAIR levels. Similar results were obtained in independent experiments shown in Fig. S2A, B. **D-G**, Immunodetection of RGA (with an antibody to RGA) in shoots (**D, E**) and roots (**F, G**) of 7-day-old wild-type (Ler) seedlings transferred from nitrate-deficient conditions to 10 mM NO_3_^-^ (**D, F**) or inversely (**E, G**), for the time indicated. Numbers represent the fold increase in RGA protein levels relative to blue-stained protein signal. Similar results were obtained in independent experiments shown in Fig. S2C-F.

### Nitrate enhances GA synthesis

An increase in the amount of GA induces a reduction in DELLA abundance, which in turn activates plant growth [41]. Moreover, GA metabolism is influenced by various environmental signals including nutrient availability [23, 26, 27]. We next examined if nitrate-deficiency-induced DELLA accumulation was attributable to a decrease in GA biosynthesis. To this end, we determined the endogenous GA contents in 7-d-old wild-type seedlings grown on low (0.5 mM) and adequate (10 mM) nitrate conditions, and 48 hr after a transfer from 0.5 to 10 mM NO_3_^-^. The levels of the major biologically active GA in *Arabidopsis*, GA_4_, as well as intermediate GAs (GA_12_ and GA_24_) were significantly increased in seedlings grown with 10 mM NO_3_^-^ compared with those of seedlings grown on low nitrate (Fig. 4A, B). Remarkably, the contents in GA_34_ (the inactive 2ß-hydroxylated product of GA_4_) were also increased in seedlings transferred to 10 mM NO_3_^-^ (Fig. 4B). This last result emphasizes that the increase in GA_4_ level is not the consequence of a reduction in the activity of the GA 2-oxidases, which catalyze the conversion of GA_4_ into GA_34_ [42].

**Figure 4.**
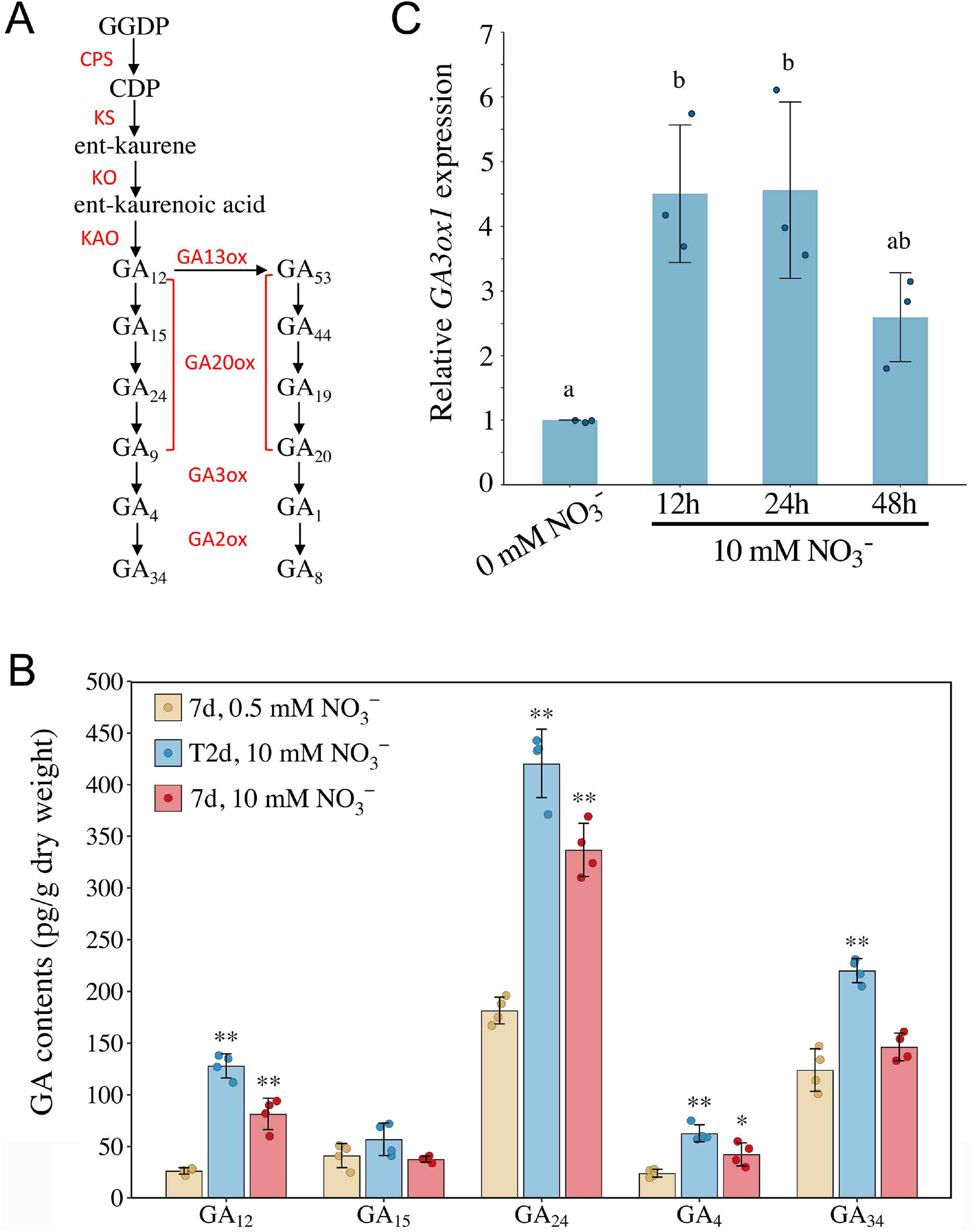
Nitrate promotes GA synthesis. **A**, Simplified GA biosynthetic pathway. Biosynthetic enzymes are highlighted in red. GGDP, geranyl geranyl diphosphate; CDP, *ent*-copalyl diphosphate; CPS, *ent*-copalyl diphosphate synthase; KS, *ent*-kaurene synthase; KO, *ent*-kaurene oxidase; KAO, *ent*-kaurenoic acid oxidase. **B**, Concentration of GAs (pg g^-1^ dry weight) in 7-day-old wild-type (Ler) seedlings grown on 0.5 or 10 mM NO_3_^-^, or two days after a transfer from 0.5 to 10 mM NO_3_^-^. Endogenous GA_9_ was analyzed but not detected. The values are means ± s.d. of four biological replicates. Asterisks indicate significant differences (\**p* < 0.05; ** *p* < 0.01) for 10 mM *versus* 0.5 mM NO_3_^-^ growth conditions by Student’s t test. **C**, Time-course of *GA3ox1* transcript accumulation in 7-day-old wild-type (Col-0) seedlings that have been transferred from 0 to 10 mM NO_3_^-^ for the time indicated. Data are means ± s.d. of three biological replicates and different letters denote significant differences (*p* < 0.05) using one-way ANOVA with Tukey’s test for multiple comparisons.

Subsequently, we investigated whether this increase in GA contents correlated with changes in the expression of GA biosynthetic genes. Surprisingly, most of GA biosynthetic genes did not show any obvious expression trend in response to a transfer from nitrate deficiency to adequate nitrate conditions, except a down-regulation for some of the genes, 48 hr after the transfer to 10 mM NO_3_^-^ (Fig. S3). However and consistent with the nitrate-induced accumulation of GA_4_ and GA_34_, we found that expression of *GA3ox1* (which catalyzes the conversion of GA_9_ into GA_4_) and *GA2ox2* is substantially up-regulated by nitrate supply (Fig. 4C and Fig. S3). When examining the expression of GA signaling genes, such as those encoding the DELLAs and GA receptors GID1, most of them did not react in response to nitrate supply, apart from a down-regulation trend 48 hr after transfer, similar to most of GA metabolism genes (Fig. S3).

### DELLA abundance is regulated by nitrate-dependent signaling

To test how nitrate controls the GA-DELLA pathway, we first examined the effects of nitrate on RGA protein accumulation in a nitrate reductase (NR)-null mutant (*nia1 nia2*; [5]). We found that nitrate decreases substantially the abundance of RGA protein in both wild-type and *nia1 nia2* mutant shoots (Fig. 5A and independent replicate shown in Fig. S4A). Thus, DELLA accumulation is regulated by nitrate signaling, independent of nitrate reduction and therefore the N status of the plant. In a subsequent experiment, we analyzed the accumulation of RGA protein in *nrt2*.*1 nrt2*.*2* (*nrt2*.*1-2*), which is mutated in two linked genes responsible for high-affinity nitrate uptake in *Arabidopsis* roots [43]. As previously reported, the *nrt2*.*1-2* double mutant displayed a reduction in shoot biomass compared with the wild-type, caused by a decrease of nitrate uptake (Fig. S4B, C). As expected, RGA accumulated to higher levels in *nrt2*.*1-2* mutant rosettes of 4-week-old plants grown on both low (0.5 mM) and non-limiting (10 mM) nitrate supply, compared with the wild-type (Fig. 5B and independent replicate shown in Fig. S4D).

**Figure 5.**
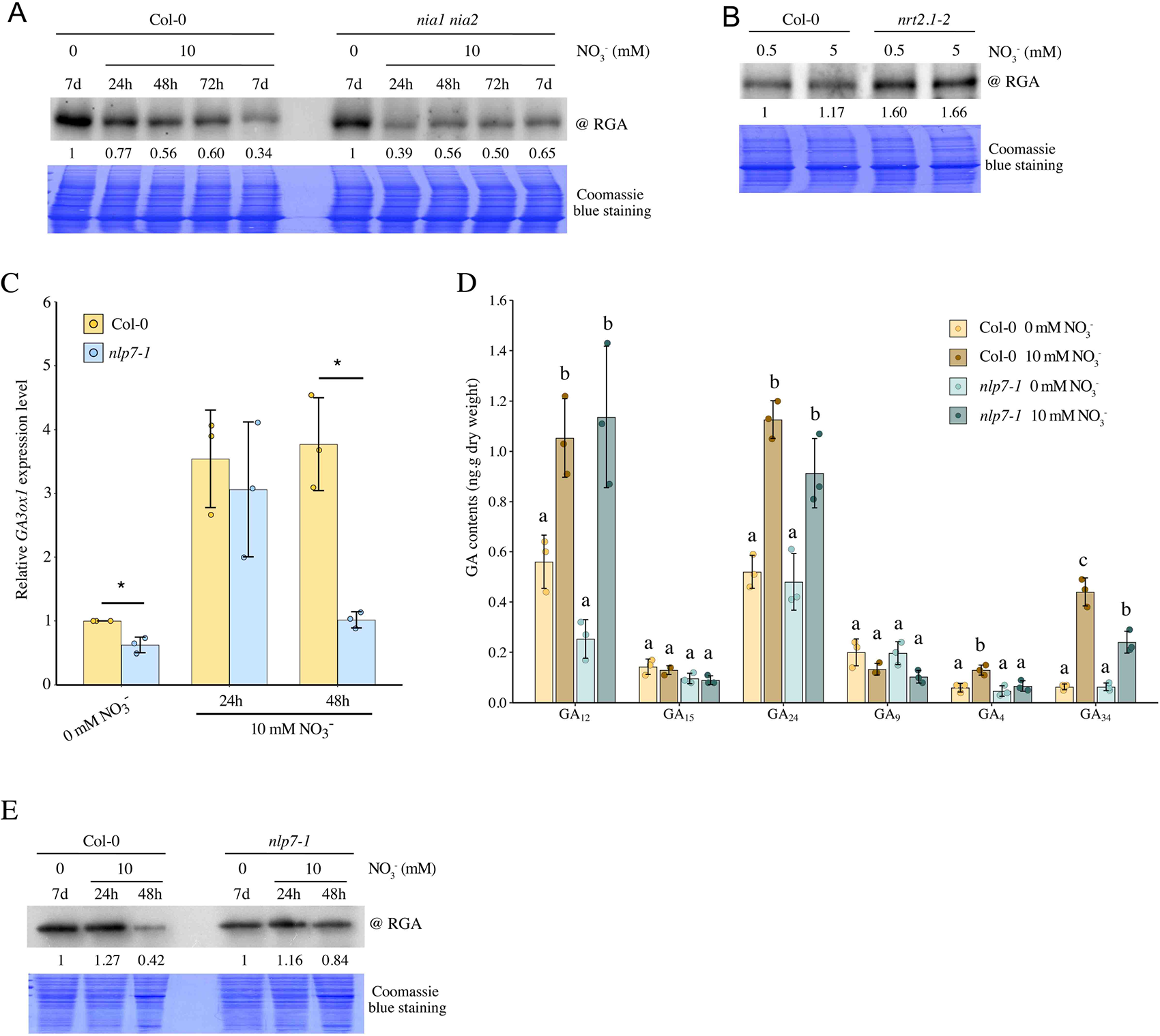
DELLA accumulation is regulated by NO_3_^-^-dependent signaling pathway. **A**, Immunodetection of RGA in shoots of 7-day-old wild-type (Col-0) and *nia1 nia2* mutant seedlings transferred from nitrate deficient conditions (0 mM) to 10 mM NO_3_^-^ for the time indicated. **B**, Immunodetection of RGA in 4-week-old wild-type (Col-0) and *nrt2*.*1-2* double mutant plants grown on 0.5 or 5 mM NO_3_^-^. **C**, Relative *GA3ox1* expression in 7-day-old wild-type (Col-0) and *nlp7-1* mutant seedlings that have been transferred from nitrate deficient conditions (0 mM) to 10 mM NO_3_^-^ for 24h or 48h. Asterisks indicate significant differences (\**p* < 0.05) for Col-0 *versus nlp7-1* by Student’s t test. **D**, GA contents (ng g^-1^ dry weight) in 7-day-old wild-type (Col-0) and *nlp7-1* mutant seedlings grown on 0 mM NO_3_^-^ or two days after a transfer from 0 to 10 mM NO_3_^-^. Different letters denote significant differences (*p* < 0.05) using two-way ANOVA with Tukey’s test for multiple comparisons. **E**, RGA protein accumulation in wild-type (Col-0) and *nlp7-1* mutant seedlings transferred from nitrate deficient conditions (0 mM) to 10 mM NO_3_^-^ for 24h or 48h. Numbers represent the fold increase in RGA protein levels relative to blue-stained protein signal. Similar results were obtained in independent experiments shown in Fig. S4A, D and E, respectively.

Key components of the nitrate-signaling pathway are the NLP6 and NLP7 transcription factors, which control the expression of a large number of early nitrate response genes [10, 14-16]. To examine the involvement of the NLP in the nitrate-regulated GA-DELLA pathway, we initially took advantage of previously reported genome-wide analyses performed on wild-type and *nlp7* mutant during a short-term nitrate resupply kinetics [10, 15]. Remarkably, these analyses revealed that even though NLP7 does not bind the genomic region of *GA3ox1* in chromatin immunoprecipitation assay, *GA3ox1* is among the list of NLP7-dependent nitrate-inducible genes [10, 15]. In agreement with these findings, we confirmed that nitrate-NLP7 pathway regulates the expression of *GA3ox1* (Fig. 5C). Whereas the expression of *GA3ox1* was up-regulated in wild-type seedlings upon nitrate supply, its expression was only transiently induced in *nlp7* mutant. Consistent with this expression pattern, nitrate supply increased the contents of bioactive GA_4_ (product of GA3ox activity) in wild-type seedlings, but not in *nlp7* mutant (Fig. 5D). By contrast, nitrate supply increased the levels of GA_12_ and GA_24_ in both wild-type and *nlp7* mutant, suggesting that nitrate regulates the early GA biosynthetic steps in an NLP7-independent manner (Fig. 5D). Finally, we analyzed the accumulation levels of RGA in wild-type and *nlp7* mutant seedlings in a time course analysis after a transfer from nitrate deficiency to adequate nitrate conditions. While nitrate supply triggered a reduction of RGA accumulation in wild-type 48 hr after transfer, the amount of RGA was relatively stable in *nlp7* mutant, in accordance with the levels of *GA3ox1* transcripts and GA_4_ contents detected in wild-type and *nlp7* mutant (Fig. 5E and independent replicate shown in Fig. S4E). Overall, these results indicate that nitrate-NLP7 signaling decreases DELLA accumulation through the activation of GA_4_ synthesis (via up-regulation of *GA3ox1* gene expression).

### Effect of nitrate on GA metabolism and growth processes in wheat

To examine whether the role of the GA-DELLA pathway in nitrate-mediated growth processes is conserved in diverse angiosperms, we analyzed the effect of nitrate (used as the only N source) on the growth of wheat seedlings (cv. Cadenza) that have altered GA signaling. We compared wild-type (*Rht-1* tall) against *Rht-D1b* (semi-dwarf) and *Rht-B1c* (severe-dwarf) gain-of-function mutants that express constitutively active DELLA (*Rht-1*) proteins conferring GA-insensitive dwarfism [20, 44]. For this experiment, wild-type, *Rht-D1b* and *Rht-B1c* mutants were grown on a perlite vermiculite soil supplemented 3 times per week with the Letcombe nutrient solution containing nitrate (+N; 1.5 mM Ca(NO_3_)_2_, 5 mM KNO_3_ and 2 mM NaNO_3_) or without N (-N; 1.5 mM CaCl_2_, 5 mM KCl, 2 mM NaCl), and the length of the primary leaf sheath was measured. As previously reported with other semi-dwarf *Rht-1* mutant alleles [17, 18], the *Rht-D1b* and *Rht-B1c* mutant wheat were relatively insensitive to nitrate supply. Whereas the length of the primary leaf sheath was substantially longer in 3-week-old wild-type plants grown with nitrate (compared with those grown without N), nitrate supply had respectively little or no effect on the primary leaf sheath length of *Rht-D1b* and *Rht-B1c* mutants (Fig. 6A, B). Thus, nitrate enhances wheat growth in part via a DELLA-dependent mechanism.

**Figure 6.**
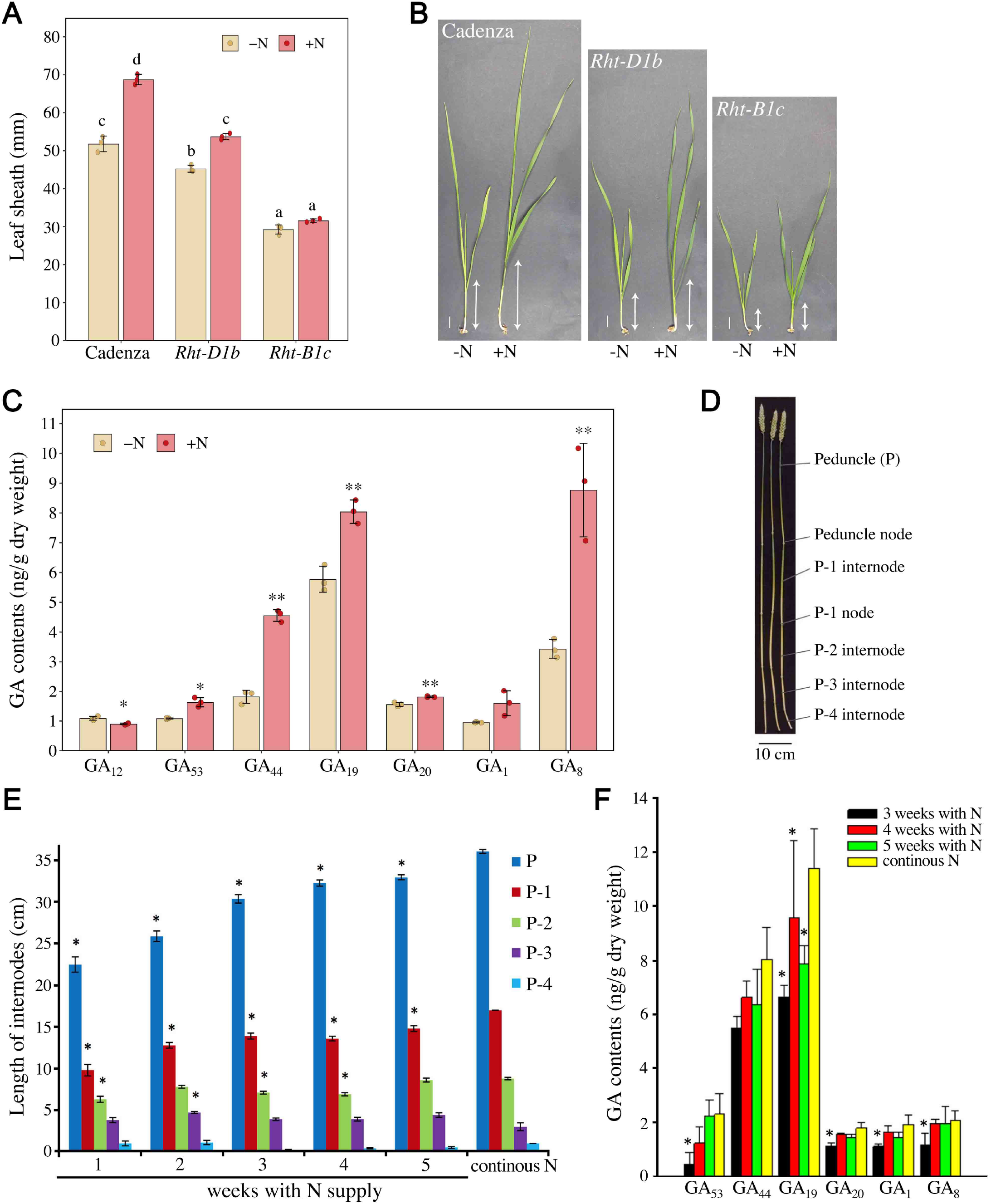
Effects of nitrate on GA metabolism and growth in wheat. **A**, Leaf sheath length (mean ± s.d. of three biological replicates) of 3-week-old wild-type (cv. Cadenza), *Rht-D1b* and *Rht-B1c* mutant wheat grown on perlite vermiculite soil supplemented 3 times per week with a nutritive solution without nitrogen (-N) or containing NO_3_^-^ (+N). Different letters denote significant differences (*p* < 0.05) using two-way ANOVA with Tukey’s test for multiple comparisons. 15 ≤ n ≤ 20 plants were analyzed for each biological replicate. **B**, Representative 3-week-old wild-type (cv. Cadenza), *Rht-D1b* and *Rht-B1c* mutant wheat grown as in **A**. The arrows indicate the length of the first leaf sheath. Scale bars represent 1 cm. **C**, Concentration of GAs (ng g^-1^ dry weight) in the first internode of 3-week-old wild-type (cv. Cadenza) wheat grown as in **A**. The values are means ± s.d. of three biological replicates. Asterisks indicate significant differences (\**p* < 0.05; ** *p* < 0.01) for +N *versus* - N by Student’s t test. **D-F**, Effect of nitrogen withdrawal on internode elongation and GA contents in wheat. (**D**) Sections of elongated stem at anthesis. P-1, second internode; P-2, third internode; P-3, P-4 further internodes. (**E**) Internode length (mean ± s.e.; n=15) of 7-week-old wild-type wheat (cv. Cadenza) growing on different nitrogen (N) status. Plants were grown on sand perlite soil supplemented three times in the first week with a nutritive solution containing NO_3_^-^ (+N). The plants were then divided into six treatment groups for which N was sequentially withdrawn from the watering solution week by week, as indicated. Asterisks indicate significant difference (*p* < 0.05) for comparisons within internodes for treatments 1-5 weeks with N compared to control, according to two-way ANOVA followed by least significant difference (LSD) test. (**F**) Concentration of GAs (ng g^-1^ dry weight) in peduncle and peduncle nodes of 7-week-old wild-type wheat grown on different N status, as in **A**. Asterisks indicate significant difference (*p* < 0.05) for comparisons within treatments (1-5 weeks with N compared to control) for each GA, according to residual maximum likelihood (REML) followed by LSD test.

As a next step, we determined the contents of GA in the first internode of 3-week-old wild-type plants grown with NO_3_^-^ or without N. In wheat, the 13-hydroxylation pathway leading to active GA_1_ is the main GA biosynthetic route [36] (Fig. 4A). We found that nitrate supply increases the levels of bioactive GA_1_, GA intermediates (GA_53_, GA_44_ and GA_19_) and inactivation product GA_8_ (Fig. 6C).

We also investigated the effect of nitrate on growth and GA synthesis at a later stage, in 7-week-old wheat cv. Cadenza (*Rht-1* tall). In this experiment, the plants were grown with nitrate and then N was sequentially withdrawn week after week, over a time scale of 6 weeks until booting stage. Noteworthy, main tillers were removed from all plants as they appeared, to avoid N resources being redistributed between tillers. In this growth context, we found that the peduncle and first two internodes were gradually shorter as time with N decreases (Fig. 6D, E). By contrast, there was no significant difference in the lengths of the third and fourth internode across treatment groups when compared to control (Fig. 6E). Consistent with these physiological traits, we found that N withdrawal had a gradual effect on endogenous GA contents in the first internode at booting stage. All GA quantified displayed a steady gradual increase with increasing length of time that the plants were supplied with nitrate (Fig. 6F). Thus altogether, and as previously shown with *Arabidopsis*, these results demonstrate that nitrate enhances wheat growth in part through activation of GA synthesis.

## Discussion

Plants are sessile organisms that must perpetually adapt their growth and development to fluctuating nitrate concentrations in the soil. It is now established that GA signaling plays a critical role in modulating plant growth in response to environmental changes including nutrient availability [22-26]. While semi-dwarfing genes that interfere with the action or production of GA contributed to substantial yield increases during the green revolution because they prevent excessive stem elongation in response to high nitrogen fertilizer regimes [17], the role of GA signaling in adapting plant growth to nitrate availability remained unclear. The experiments described here investigate the connections between the GA pathway and the nitrate-dependent signaling pathway.

Plant root development is strongly influenced by nitrate concentrations in the soil. Whereas limited nitrogen provisions inhibit the primary root length, excess supply of nitrate also leads to a repression of root elongation [32, 45]. Remarkably, we observed that low (0.5 mM NO_3_^-^) and high (50 mM NO_3_^-^) nitrate provisions promoted the accumulation of RGA in roots and that the growth of *global-DELLA* mutant roots was less affected by these extreme nitrate concentrations than those of wild-type seedlings. Furthermore, the abundance of RGA in wild-type roots and the difference in root growth between wild-type and *global-DELLA* mutant seedlings decreased progressively with increase in nitrate supply, from low (0.5 mM NO_3_^-^) to adequate (10 mM NO_3_^-^) concentrations. Thus, low (and high) nitrate conditions repress root growth in part via a DELLA-dependent pathway. Consistent with our findings, a previous work has reported a similar effect of high nitrate supply on plant growth and GFP-RGA accumulation in *pRGA:GFP-RGA* seedlings roots, except that the range of NO_3_^-^ concentrations used by the authors was ten-fold less than in our growth conditions [46]. This difference in effective NO_3_^-^ levels could be attributed to the composition of the N-free medium, which is different between the two works. Interestingly, we also found that seedling shoots had a similar response to roots in terms of RGA accumulation, even though the shoots tend to be less sensitive to nitrate than roots. Consistent with this observation, global gene expression analyses revealed that while roots treated with low nitrate have a much wider response than shoots in term of the number of genes altered, the shoots can be as responsive as roots to adequate nitrate concentrations [5, 47]. Thus, it is likely that in our growth conditions, 1 mM NO_3_^-^ supply is the minimal concentration to trigger a decrease in RGA abundance in seedling roots, while 10 mM NO_3_^-^ is required to observe the same phenomenon in shoots. Moreover, the kinetics of DELLA disappearance in response to a transfer from nitrate deficiency to adequate nitrate conditions (and reciprocally) is similar in both roots and shoots, and is visible within a few days after transfer.

Our results also demonstrate that nitrate promotes stem elongation and shoot branching in *Arabidopsis* and leaf sheath elongation in wheat in part through a DELLA-dependent mechanism. Thus nitrate-mediated regulation of DELLA accumulation is a general mechanism that is conserved among flowering plants and throughout plant development. Consistent with this finding, we found that nitrate enhances the accumulation of bioactive GAs in both *Arabidopsis* and wheat. At this stage, it remains unclear whether nitrate enhances the accumulation of bioactive GA_1_ in wheat through the NLP-dependent signaling pathway, such as in *Arabidopsis*. Interestingly, not only the bioactive GAs increased in response to nitrate supply, but also GA_12_ and GA_53_ in *Arabidopsis* and wheat, respectively, and later intermediate GAs. This implies increased activity of the earlier part of the GA biosynthetic pathway and enhanced metabolic flux through the pathway (Fig. 4A). Overexpression of *ent-*copalyl diphosphate synthase (CPS) and *ent*-kaurene synthase (KS) in *Arabidopsis* massively increases the supply of GA_12_ but this has almost no effect on the levels of bioactive GAs [48]. Thus, although it is likely that nitrate regulates the GA pathway at multiple levels, it enhances the production of bioactive GAs via its effect on the final steps of GA biosynthesis. It is worth noting that while our results in *Arabidopsis* indicate that nitrate induces the expression of *GA3ox1*, we cannot exclude the possibility of post-translational regulation of GA biosynthetic enzymes (e.g. protein turnover) and allosteric control of enzyme activity, two important mechanisms largely understudied [49, 50].

Previous works revealed that bioactive GAs promote root growth through cell division and expansion by stimulating the degradation of DELLAs in endodermal cells of the RAM and root elongation zone, respectively [38-40]. In elongating roots, bioactive GAs are translocated in the endodermis by the GA transporter NITRATE TRANSPORTER1/PEPTIDE TRANSPORTER FAMILY 3 (NPF3) localized in the plasma membrane of endodermal cells [51, 52]. We also observed that adequate nitrate supply enhances primary root growth by activating both cell proliferation and elongation. However, unexpectedly, whereas nitrate increased the number of cortical cells in root meristem via a DELLA-dependent mechanism, it increased cell elongation in a DELLA-independent manner. DELLAs restrain cell cycle activity by enhancing the expression of cell cycle inhibitors [40]. Accordingly, we found that mitotic activity (monitored with the CYCB1;2:Dbox-GUS reporter) was less reduced in the RAM of the DELLA loss of function *gai-t6 rga-24* mutant grown on low nitrate conditions than those of wild-type. By contrast, low nitrate restrained cell length of both wild-type and *global-DELLA* mutant seedling roots. Thus, it is likely that nitrate-induction of cell size is driven by other hormonal pathways [31]. Of note, it has recently been reported that nitrate promotes cell size by increasing ploidy level [53].

It is now well established that the nitrate-signaling pathway activates plant growth by controlling the hormonal status of the plant [30, 31]. Reciprocally, hormone-signaling pathways regulate nitrate acquisition and assimilation, constituting a retro-control of growth on N provision. A recent report revealed that *Rht-1* dwarfing genes restrain nitrate uptake and assimilation by interacting with and inhibiting the activity of GRF4, a transcription factor that positively regulates N metabolism [29]. We propose that GA-nitrate interplay provides a highly regulated mechanism permitting flexible and appropriate growth in response to N availability.

### Lead contact and materials availability

All plant lines are available for sharing. Further information for resources and reagents should be forwarded and attended by the Lead Contact, Patrick Achard.

### Experimental Model and Subject Details

#### Plants

Mutants and transgenic lines in *Arabidopsis* were derived from Landsberg *erecta* (*global DELLA*; *ga20ox1* (formerly *ga5*); *gai-t6 rga-24*; *pRGA:GFP-RGA*) or Columbia-0 (*nia1 nia2*; *nrt2*.*1-2*; *nlp7-1*; *pCYCB1,2:Dbox-GUS*) ecotypes. To generate *gai-t6 rga-24 pCYCB1,2:Dbox-GUS*, the *pCYCB1,2:Dbox-GUS* (Col-0 ecotype) was crossed to *gai-t6 rga-24* (Ler background) to obtain an F1 population. The F3 plant was selected and then backcrossed one time into *pCYCB1,2:Dbox-GUS* Col-0 genetic background. Two independent *gai-t6 rga-24 pCYCB1,2:Dbox-GUS* homozygous lines (#7 and #18) were used in the study. Wheat near isogenic lines (cv Cadenza) containing *Rht-D1b* and dwarf *Rht-B1c* alleles were used in this study. The *Rht-B1c* allele was introduced from cultivar Mercia [55]. The *Rht-D1b* allele was transferred from cultivar Avalon [54]. The alleles were selected from homozygous progenies after six backcrosses to cultivar Cadenza (which contains the wild-type *Rht-1* alleles: *Rht-A1a, Rht-B1a* and *Rht-D1a*) with recurrent selection for the dwarfing mutation.

### Method details

#### Growth conditions for Arabidopsis

*In vitro*, plants were grown on 1x Murashige-Skoog (MS) modified medium without nitrogen (Ref 30630200; bioWORLD plant media) supplemented with an adequate KNO_3_ concentration or 0.5 mM glutamine (as indicated in the figures), 0.5 mM NH_4_-succinate, 1% sucrose and 1% ultrapure agar (Merck) or 1% type A agar (Sigma) under a 16 h photoperiod at 22°C, except for the analysis of the expression of GA metabolism genes, for which the plants were grown in continuous light. In soil, plants were grown in pots on a mix perlite/vermiculite (1/2 v/v) completed with 5% soil under long day photoperiod (16h light at 21°C / 8h dark at 18°C). The pots were watered once a week with 5 ml of 1x MS modified medium without N supplemented with KNO_3_ as indicated (K^+^ were adjusted with KCl).

#### Growth conditions for wheat

*Rht-1* tall, *Rht-B1c* and *Rht-D1b* plants were grown on a mix perlite/vermiculite (1/2 v/v) supplemented with 5% soil under long day photoperiod (16h light at 20°C / 8h dark at 15°C). The pots were watered three times a week with 13 ml of 1x Letcombe modified nutrient solution without N (N-deficiency condition) or with nitrate (N+ condition; 1.5 mM Ca(NO_3_)_2_, 5 mM KNO_3_ and 2 mM NaNO_3_). The composition of Letcombe modified nutrient solution is listed in Table S1. For N withdrawal experiment, the seeds were sown on sand/perlite pots and watered three times per week.

The first week, all plants were watered with N+ solution, then with either N+ or N-solution. N was sequentially withdrawn week after week over a time scale of 6 weeks until booting stage.

#### Immunoblot analyses

Seedling roots or shoots were ground in 2x SDS-PAGE sample buffer followed by boiling for 5 min. After centrifugation, the protein extracts were fractionated on a 8% or 10% SDS-PAGE gel and blotted onto membrane. Immunoblots were performed using a 2000-fold dilution of anti-RGA (Agrisera) or anti-GFP (Clontech), and a 5000-fold dilution of peroxidase-conjugated goat anti-rabbit or mouse IgG (Molecular Probes). Signals were detected with Fusion FX (Vilber) using the Luminata Forte Western HRP Substrate (Millipore) or Clarty Max™ Western ECL substrate (Biorad). The blot was subsequently probed with anti-cdc2 (PSTAIRE) antibody (Santa Cruz Biotechnology) for loading control or stained with Coomassie blue. Quantification of the signals was performed using ImageJ package version 1.48v (www.imageJ.nih.gov).

#### Gene expression analyses

Total RNA was extracted using RNeasy Plant mini kit (Qiagen) according to manufacturer’s instructions. qRT-PCR was performed using gene-specific primers (listed in Table S2) on a Lightcycler LC480 apparatus (Roche). AT4G26410 gene was used as internal reference transcript. The relative expression level of each gene was calculated using Lightcycler 480 software, release 1.5.0 SP3.

#### Observation of GFP fluorescence

GFP-RGA fluorescence was determined on 7 day-old *pRGA:GFP-RGA* root tip, with a Zeiss LSM780 inverted confocal laser microscope with x20 objective. All images were obtained with the same modifications and intensity parameters. Root cell walls were stained with propidium iodide (10 µg/mL, Sigma).

#### GUS analyses

Histochemical detection of GUS activity was carried out on roots of 5-d-old *pCYCB1*.*2:Dbox-GUS* and *gai-t6 rga-24 pCYCB1*.*2:Dbox-GUS* seedlings grown in N-deficient or nitrate conditions as indicated. Plant material was infiltrated in GUS solution (250 µg/ml 5-bromo-4-chloro-3-indolyl-*ß*-D-glucuronide (X-Gluc); 50 mM sodium phosphate pH 7; 1 mM potassium ferricyanide; 1 mM potassium ferrocyanide; 10 mM EDTA; 0.01% triton X-100) for 15 min and incubated at 37°C overnight. Then, the GUS solution was replaced with 100% (v/v) ethanol during 6 h at room temperature and kept in 70% (v/v) ethanol at 4°C. For root observations, 70% (v/v) ethanol was replaced with 0,24N HCl in 20% methanol during 15 min at 57°C. Then, this solution was replaced by 7% NaOH in 60% ethanol during 15 min at room temperature. Finally, seedlings were rehydrated with successive bath of 40%, 20% and 10% ethanol during 5 min each and kept in 5% ethanol 25% glycerol. Roots were observed in 50% glycerol using AxioImager Z1 (Zeiss) microscope.

#### Root length, root meristem size and cell length analyses

Seeds of wild-type (Ler) and *global-DELLA* mutant were plated on 1x MS modified medium without nitrogen supplemented with nitrate at the concentration indicated, and placed at 4°C at least 2 days to synchronize germination. Plates were then placed vertically in a growth chamber (22°C; 16h photoperiod). As *global-DELLA* mutant seeds germinate earlier than wild-type, the plates containing wild-type seeds were disposed in the growth chamber 8h before those containing *global-DELLA* mutant seeds. Seeds that have germinated synchronously (protrusion of the radicle) were marked (the other seeds were discarded) and root length was measured 7 days post-germination (DPG). Meristem size was expressed as the number of cells in cortex files from the QC to the first elongated cell exhibiting vacuolization. Measurements were performed at 3, 5 and 7 DPG by microscopy (AxioImager Z1, Zeiss) on at least 50 roots for each genotype. Length of differentiated root cortical cells was measured with Image J software (http://rsb.info.nih.gov).

#### GA determinations

GA contents in *Arabidopsis* seedlings grown in liquid MS modified medium without nitrogen and supplemented with indicated nitrate concentrations (Fig. 4B and 5D) and in the primary internode of 3-week-old wheat grown perlite/vermiculite supplemented with Letcombe modified nutrient solution (Fig. 6C) were determined by ultra-high performance liquid chromatography-mass spectrometry (UHPLC-MS) using a Q-Exactive spectrometer (Orbitrap detector; ThermoFisher Scientific). Dry grounded material was suspended in 80% methanol-1% acetic acid including 17-^2^H_2_-labeled GA internal standards (Olchemim) and mixed by shaking during one hour at 4°C. The extract was kept at -20°C overnight, centrifuged and the supernatant dried in a vacuum evaporator. The dry residue was dissolved in 1% acetic acid and passed through an Oasis HLB column. The dried eluate was dissolved in 5% acetonitrile-1% acetic acid, and the GAs were separated by UHPLC (Accucore RP-MS column 2.6 µm, 50 × 2.1 mm; ThermoFisher Scientific) with a 5 to 50% acetonitrile gradient containing 0.05% acetic acid, at 400 µL/min over 14 min. The concentrations of GAs in the extracts were analyzed by selected ion monitoring (SIM) using embedded calibration curves and the Xcalibur 2.2 SP1 build 48 and TraceFinder programs.

GA contents in peduncle and peduncle node of 7-week-old wheat grown on different N status (Fig. 6F) were determined by gas chromatography-mass spectrometry with selected ion monitoring (SIM) of endogenous and 17-^2^H_2_-labeled GA internal standards as described previously [55].

### Quantification and Statistical Analysis

#### Root measurements

Root length and length of differentiated root cortical cells were measured with Image J software (http://rsb.info.nih.gov).

#### Statistical tests

All graphs, box plots and histograms, as well as statistical analyses were generated with RStudio package v.1.2.1335 (www.rstudio.com). Phenotypic characterization, quantitative PCR with reverse transcription in wheat and GA quantifications were analyzed using Student’s *t*-test (*p <* 0.01 or 0.05) or by analysis of variance (ANOVA). Tukey’s Honest Significant Difference (HSD) was used to compare between genotypes and nitrogen conditions using a significance threshold of 5%. Statistical analyses of the figure 6E, F were performed using the GenStat statistical software (14th edition, 2011, VSN International Ltd., Hemel Hempstead, UK).

## Data and Code Availability

This work does not involve the production of large datasets and uses published plugins or image analysis tools.

## Acknowledgments

We thank Tp. Sun for providing seeds of *pRGA:GFP-RGA*, J. Leon for *nia1 nia2* mutant and the NASC for *pCYCB1;2:Dbox-GUS* and *nrt2*.*1-2* mutant. This work was supported by the Centre National de la Recherche Scientifique and the French ministry of research and higher education. B.G. was supported by a studentship from the Lawes Agricultural Trust. A.P. and S.T. were supported by Institute Strategic Programme grant BBS/E/C/000I0220 from the Biotechnology and Biological Sciences Research Council of the UK. PH was supported by The Czech Science Foundation (grant Nos 18-10349S and 20-17984S) and the European Regional Developmental Fund Project “Centre for Experimental Plant Biology” No. CZ.02.1.01/0.0/0.0/16_019/0000738.

## Author contributions

All the experimental work, except GA quantifications and figure 6E, F, was performed by L.C., L.J., L.S.A, J.M.D and P.A. GA quantifications were performed by B.G., E.C., J.K., D.H., S.G.T., P.H. and A.L.P. Experimental work shown in figure 6E, F was performed by B.G., S.G.T., P.H. and A.L.P. L.C., L.J., M.S.J., P.A., J.M.D, L.S.A, B.G., S.T., P.H., G.K., S.R. and A.L.P. designed the experiments and analyzed the results; P.A., L.C., J.M.D, G.K., S.R., S.G.T., P.H. and A.L.P. wrote the paper.

## Supplemental Data

### Supplemental Figures and Tables

**Figure S1.**
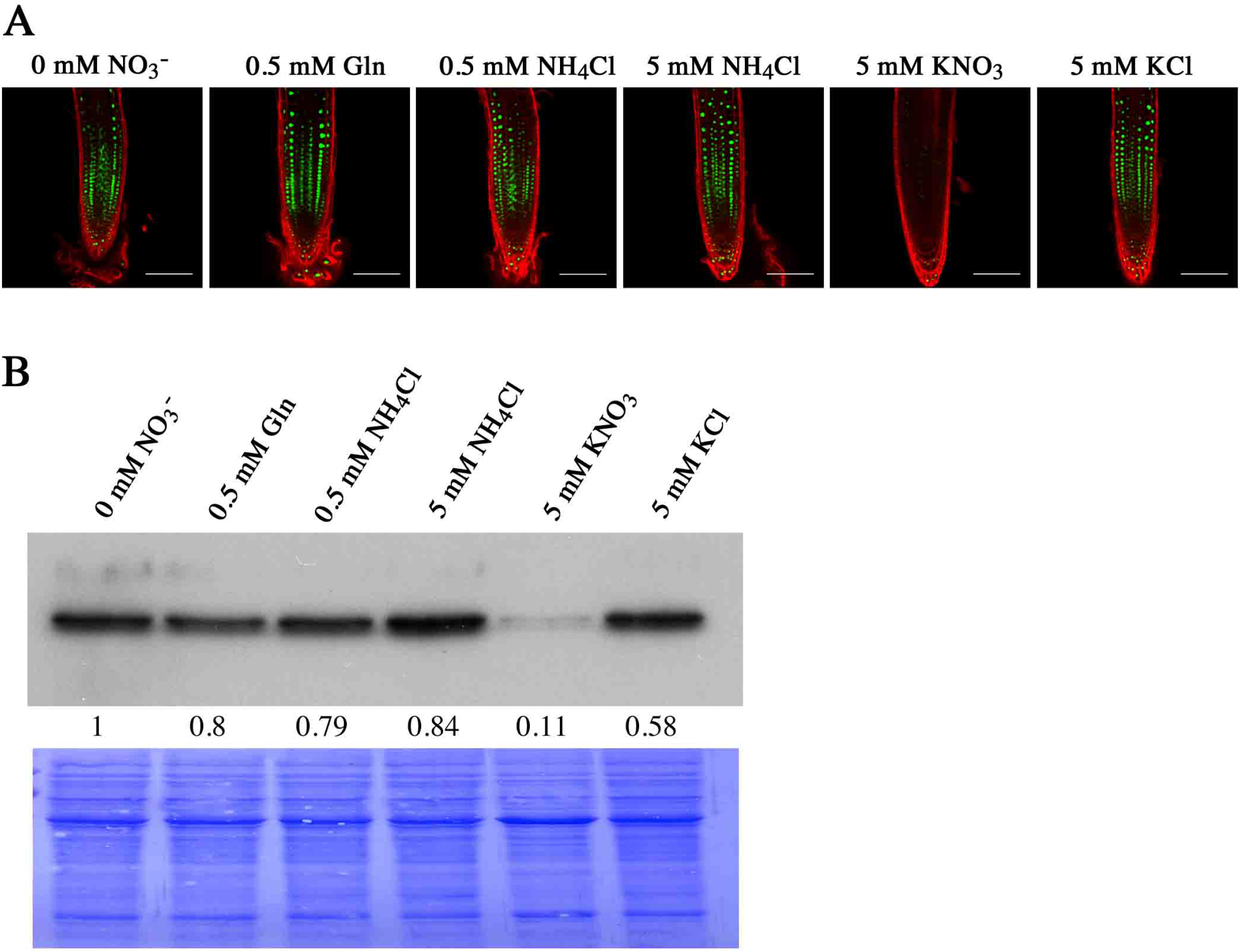
DELLA accumulation is dependent on nitrate availability. **A**, GFP fluorescence in root meristem of 7-day-old *pRGA:GFP-RGA* seedlings grown on media without nitrate or complemented with 0.5 mM Gln, 0.5 mM or 5 mM NH_4_Cl, 5 mM KNO_3_, or 5 mM KCl, as indicated. All images were obtained with the same parameters. Scale bars represent 100 µm. **B**, Immunodetection of GFP-RGA (with an antibody to GFP) in roots of 7-day-old *pRGA:GFP-RGA* seedlings grown in same conditions as in **A**. Numbers represent the fold increase in GFP-RGA protein levels relative to blue-stained protein signal.

**Figure S2.**
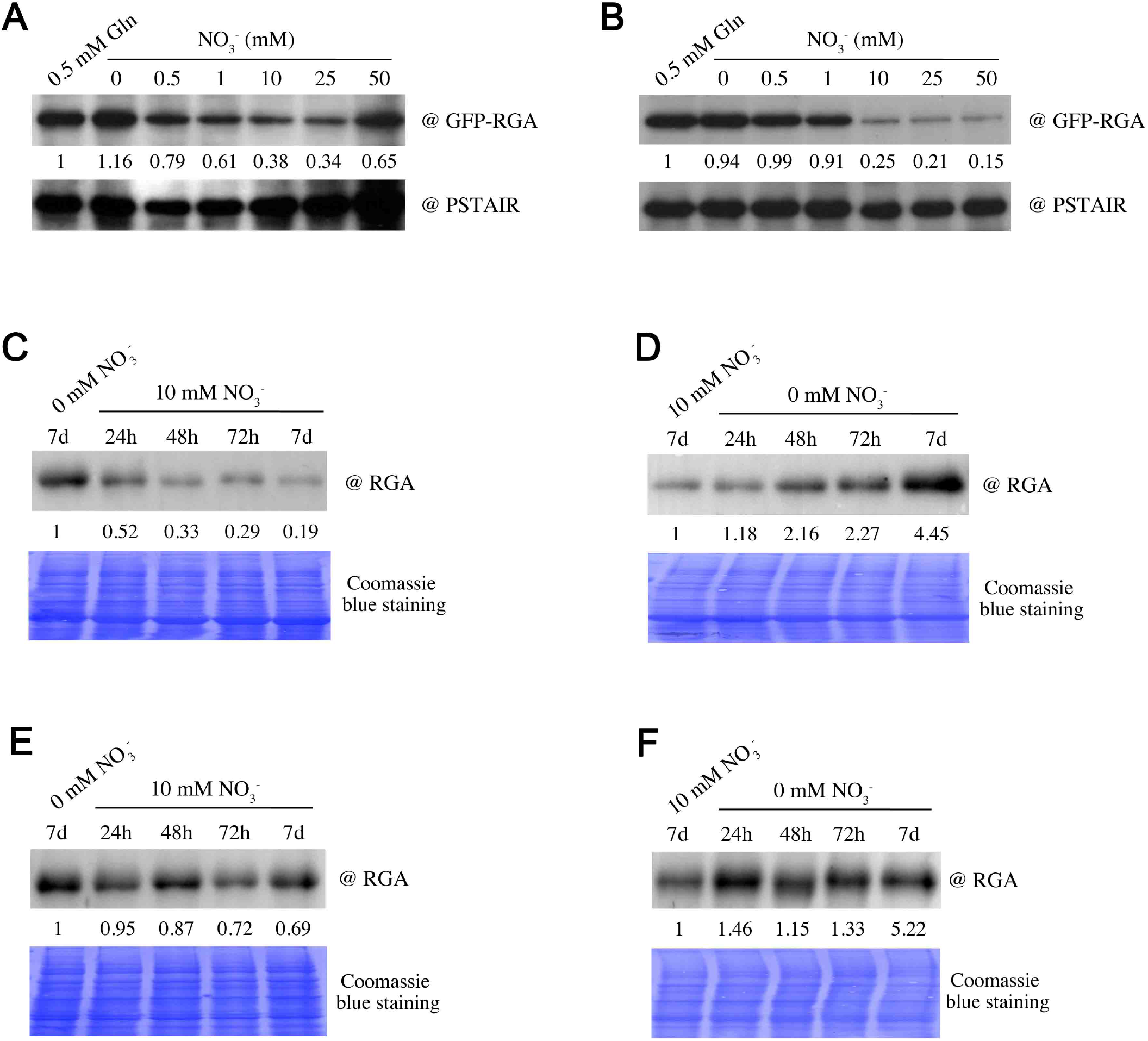
Nitrate enhances DELLA protein degradation. **A-B**, Immunodetection of GFP-RGA in roots (**A**) and shoots (**B**) of 7-day-old *pRGA:GFP-RGA* seedlings grown on media containing increasing concentration of NO_3_^-^, as indicated. PSTAIR serves as sample loading control. Numbers represent the fold increase in GFP-RGA protein levels relative to PSTAIR levels. Biological replicates of Fig. 3B and C. **C-F**, Immunodetection of RGA in shoots (**C, D**) and roots (**E, F**) of 7-day-old wild-type (Ler) seedlings transferred from nitrate-deficient conditions to 10 mM NO_3_^-^ (**C, E**) or inversely (**D, F**), for the time indicated. Numbers represent the fold increase in RGA protein levels relative to blue-stained protein signal. Biological replicates of Fig. 3D-G.

**Figure S3.**
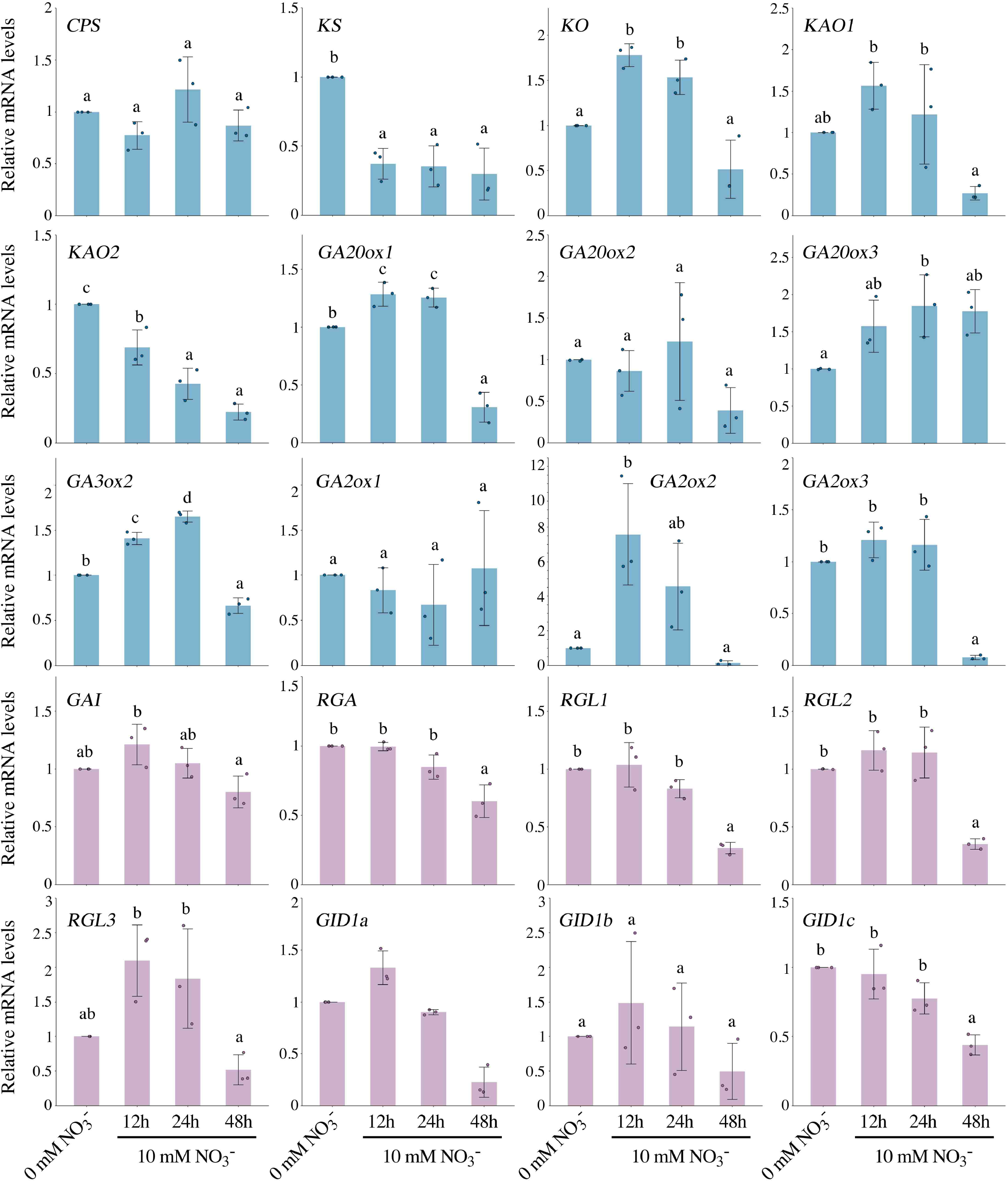
Effect of nitrate on GA metabolism and signaling gene expression. Expression levels of selected GA biosynthetic and signaling genes in 7-day-old wild-type (Col-0) seedlings transferred from 0 to 10 mM NO_3_^-^ as indicated. Data are means ± s.d. of three biological replicates and different letters denote significant differences (*p* < 0.05) using one-way ANOVA with Tukey’s test for multiple comparisons.

**Figure S4.**
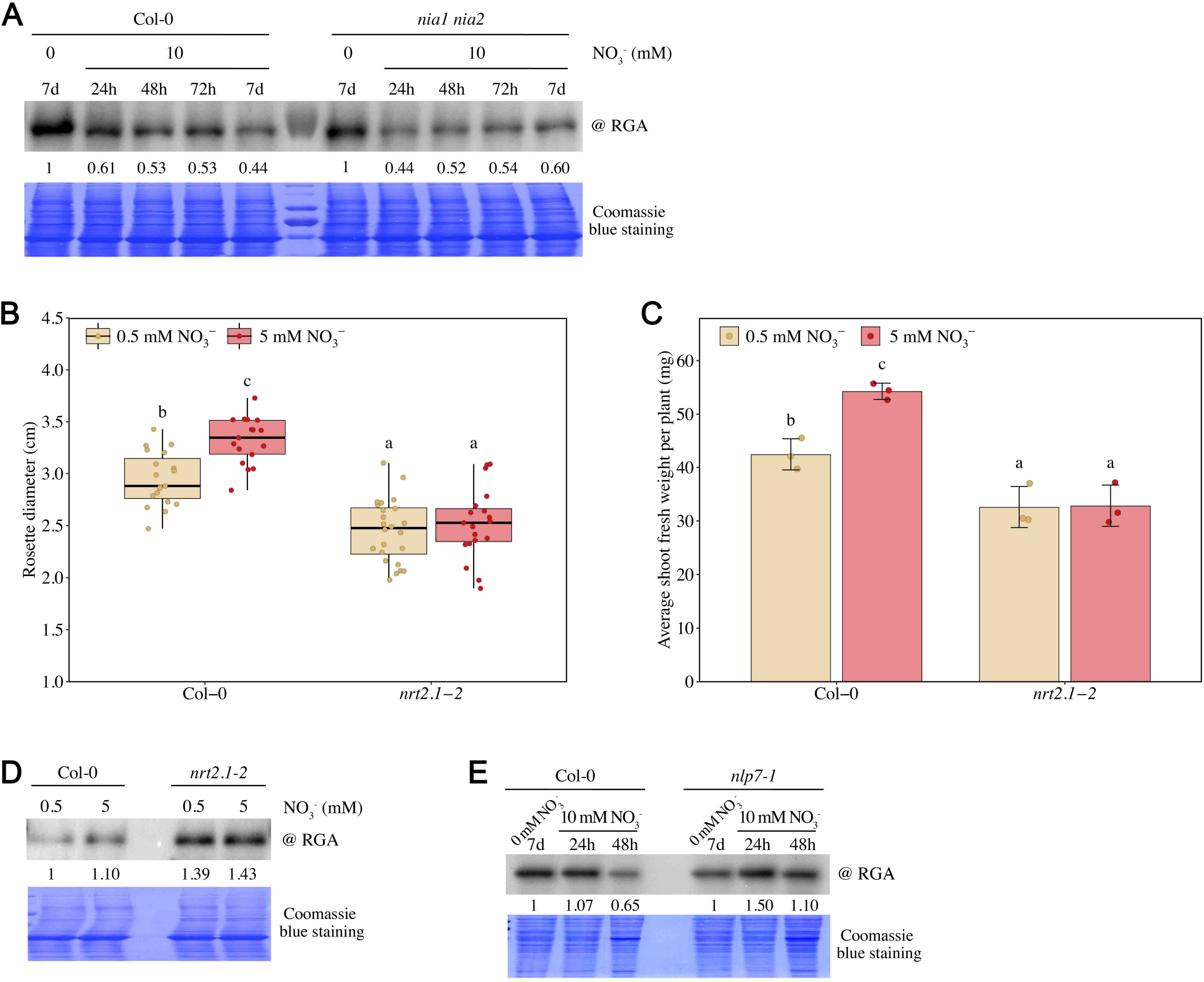
GA-DELLA pathway is regulated by nitrate-dependent signaling. **A**, Immunodetection of RGA in shoots of 7-day-old wild-type (Col-0) and *nia1 nia2* mutant seedlings transferred from nitrate deficient conditions (0 mM) to 10 mM NO_3_^-^ for the time indicated. Biological replicate of Fig. 5A. **B-C**, Rosette diameter (mean ± s.d. of 17 ≤ n ≤ 24 plants) (**B**) and shoot fresh weight (mean ± s.d. of three biological replicates; each replicate containing at least 14 ≤ n ≤ 24 plants) (**C**) of 4-week-old wild-type (Col-0) and *nrt2*.*1-2* double mutant grown on 0.5 or 5 mM NO_3_^-^. Different letters denote significant differences (*p* < 0.05) using two-way ANOVA with Tukey’s test for multiple comparisons. **D**, Immunodetection of RGA in 4-week-old wild-type (Col-0) and *nrt2*.*1-2* double mutant plants grown on 0.5 or 5 mM NO_3_^-^. Biological replicate of Fig. 5B. **E**, Immunodetection of RGA in 7-day-old wild-type (Col-0) and *nlp7-1* mutant seedlings transferred from nitrate deficient conditions (0 mM) to 10 mM NO_3_^-^ for the time indicated. Biological replicate of Fig. 5E.

**Table S1.**
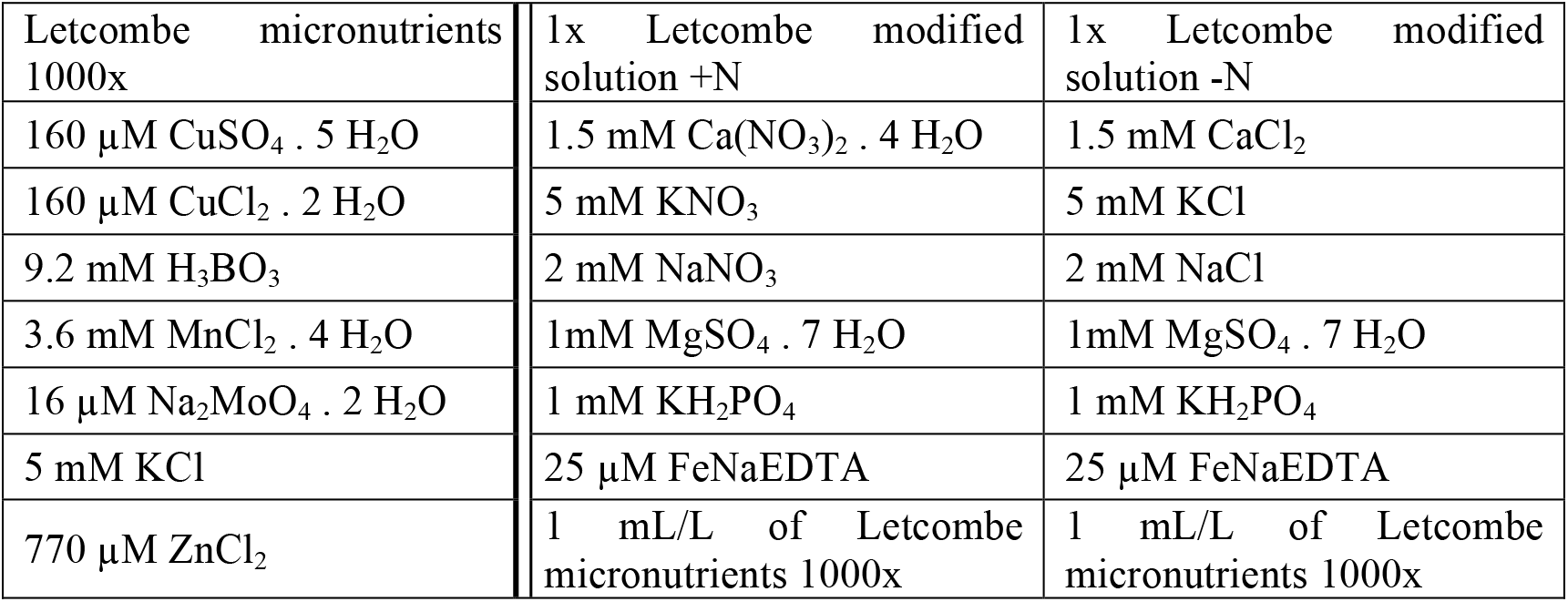
Letcombe modified solution for wheat.

**Table S2.**
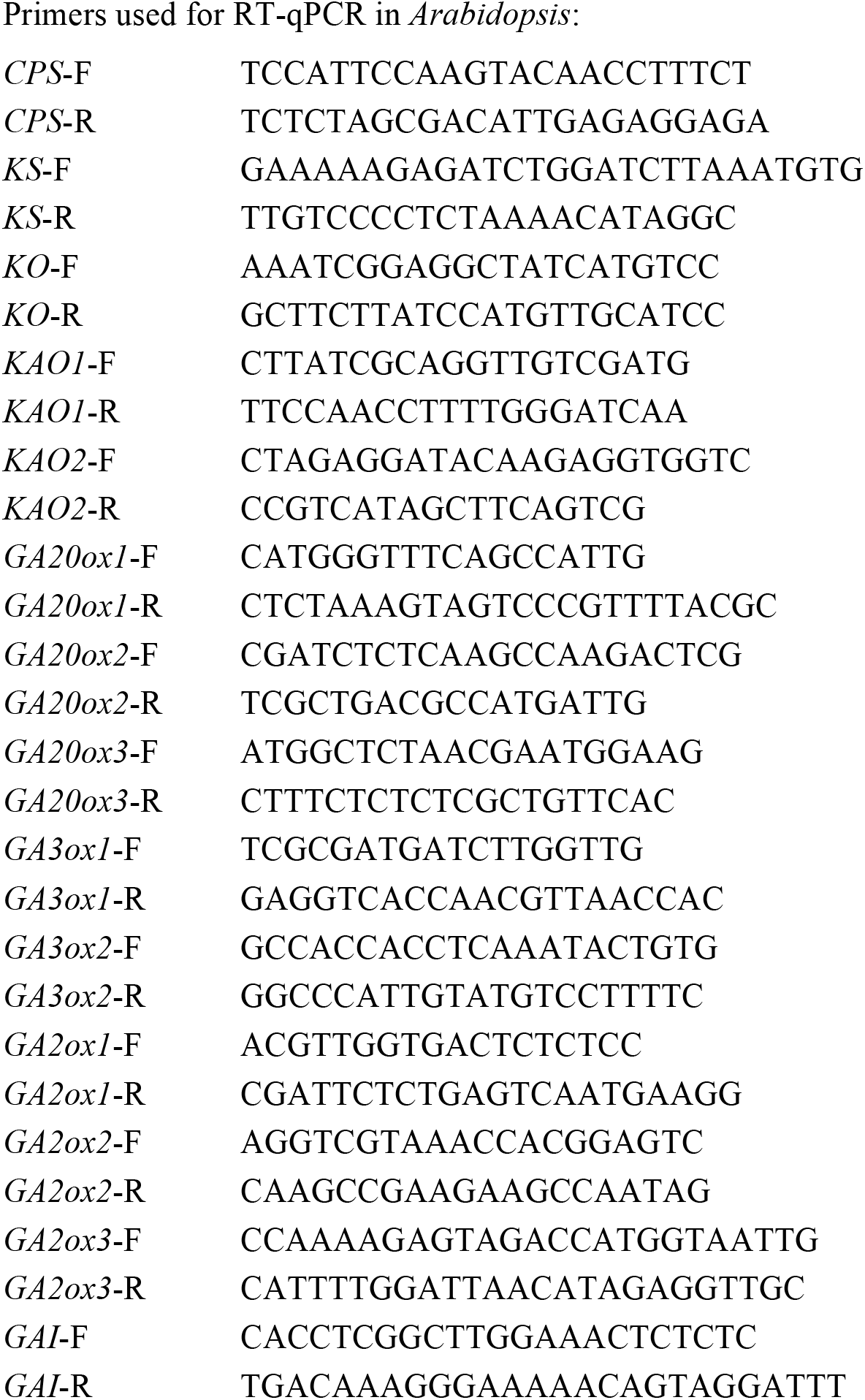
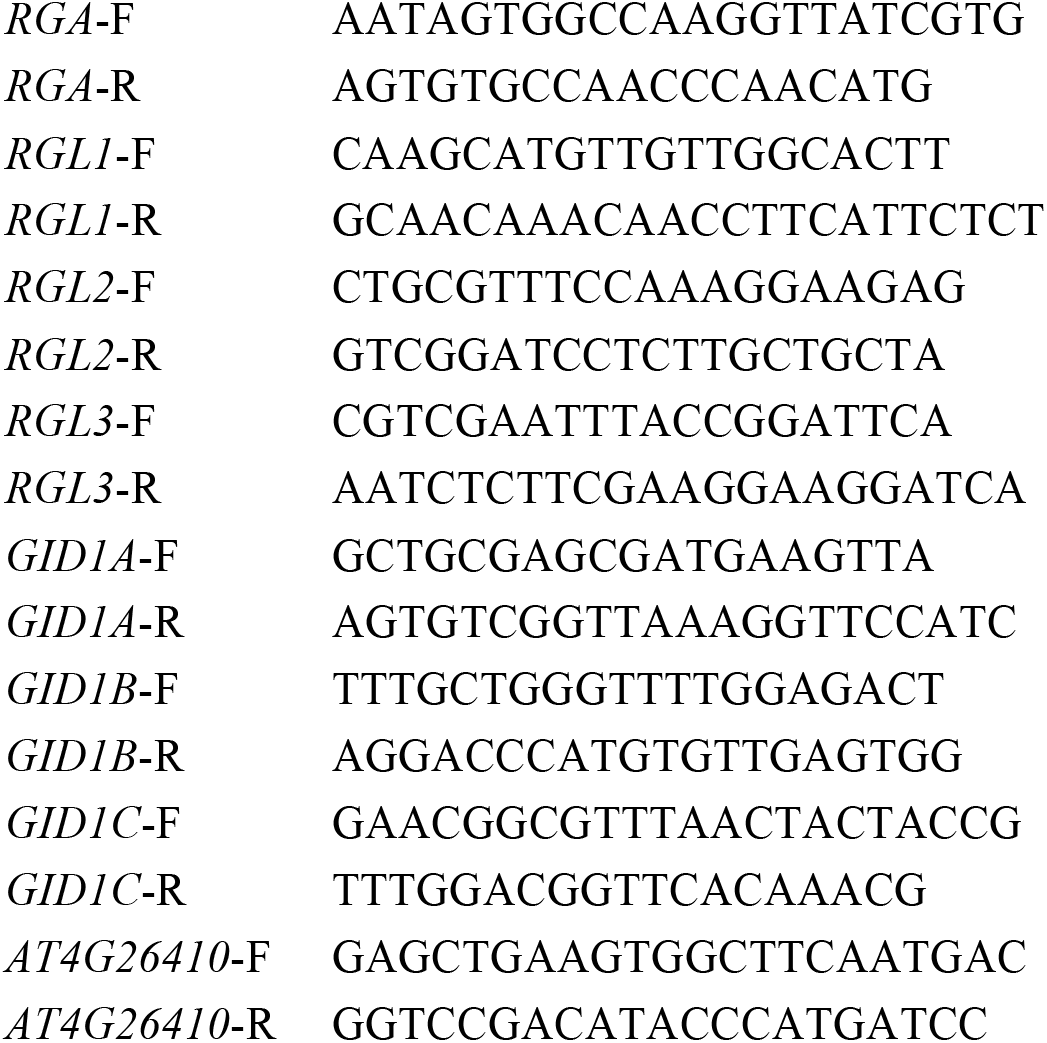
List of primers used in the study.

